# Estimating Autoantibody Signatures To Detect Autoimmune Disease Patient Subsets

**DOI:** 10.1101/128199

**Authors:** Zhenke Wu, Livia Casciola-Rosen, Ami A. Shah, Antony Rosen, Scott L. Zeger

## Abstract

Autoimmune diseases are characterized by highly specific immune responses against molecules in self-tissues. Different autoimmune diseases are characterized by distinct immune responses, making autoantibodies useful for diagnosis and prediction. In many diseases, the targets of autoantibodies are incompletely defined. Although the technologies for autoantibody discovery have advanced dramatically over the past decade, each of these techniques generates hundreds of possibilities, which are onerous and expensive to validate. We set out to establish a method to greatly simplify autoantibody discovery, using a pre-filtering step to define subgroups with similar specificities based on migration of radiolabeled, immunoprecipitated proteins on sodium dodecyl sulfate (SDS) gels and autoradiography [**G**el **E**lectrophoresis and band detection on **A**utoradiograms (GEA)]. Human recognition of patterns is not optimal when the patterns are complex or scattered across many samples. Multiple sources of errors - including irrelevant intensity differences and warping of gels - have challenged automation of pattern discovery from autoradiograms.

In this paper, we address these limitations using a Bayesian hierarchical model with shrinkage priors for pattern alignment and spatial dewarping. The Bayesian model combines information from multiple gel sets and corrects spatial warping for coherent estimation of autoantibody signatures defined by presence or absence of a grid of landmark proteins. We show the pre-processing creates more clearly separated clusters and improves the accuracy of autoantibody subset detection via hierarchical clustering. Finally, we demonstrate the utility of the proposed methods with GEA data from scleroderma patients.

## 1. Introduction

Discovering disease subgroups that share distinct disease mechanisms is fundamental to disease prevention, monitoring and treatment. For example, in autoimmune diseases, specific autoimmune responses are associated with distinct disease phenotypes and trajectories (Rosen and Casciola-Rosen, 2016). Defining the molecular markers of these subgroups has value, as these markers are of diagnostic and prognostic significance, and guide management and therapy. For example, an immune response to RNA polymerase III in scleroderma is associated with cancer; this immune response arises in response to a mutation in RNA polymerase III in that patient’s cancer. While many prominent specificities recognized by the immune response have been defined, many remain to be discovered. Although modern measurement technologies are revolutionizing the ability to define specificities, each technique results in hundreds of possibilities, which are onerous and expensive to validate. A simple technique identifies patterns of antibody reactivity based on the abundance of different weighted antigens immunoprecipitated by patient sera. Defining similar antibody reactivity patterns prior to applying one of the new discovery technologies would greatly simplify validation and therefore reduce the cost and improve the speed of antigen identification.

To identify the autoantibodies present in a patient’s serum, scientists mix serum collected from each patient with radiolabeled lysates made from cultured cells. These lysates contain a representation of all the proteins expressed in that cell type. Antibodies in each patients serum recognize and bind tightly to the specific protein(s) in the lysate against which they are directed (termed immunoprecipitation). After further processing, electrophoresis is used to sort the immunoprecipitated mixture of molecules using a crosslinked polymer or gel that separates the proteins by weight. Because different weighted molecules migrate with different speeds, the sorted molecules form distinct autoradiographed bands along the gel. By design, one gel can sort multiple samples on parallel lanes. Such experiments, referred to as gel electrophoresis autoradiography (GEA), serve to identify subsets that share one or more interesting observed bands. It is noteworthy that the lysate proteins are present in their native conformation. In our experience, many autoantibodies have epitopes that are conformationally dependent, giving GEA a powerful advantage over many of the new peptide-based (linear epitopes) sequencing technologies. The method in this paper is designed to estimate a multivariate binary autoantibody signature for each sample that represents the presence or absence of autoantibodies over a grid of molecular weights, referred to as *landmarks*.

To infer patient subsets, we can cluster patients based upon the presence or absence of each band as well as other features of the radioactive intensities such as the peak scale and amplitude. There are two critical barriers to the successful implementation of this approach that we address. First, there are *batch*, or *gel effects* in the raw GEA data. By design, molecules of identical weight would migrate the same distance along the gel. This distance however varies by gel due to differential experimental conditions. Second, gels are frequently slightly warped as they electrophorese due to heating effects generated during the electrophoresis procedure and due to artifacts introduced during physical processing of the gels. As the size and complexity of GEA experiment database grow, the need for systematic, reproducible and scalable error correction has also grown.

In this paper, we introduce and illustrate a novel statistical approach to pre-process the high-frequency GEA data which we show improve our ability to compare and cluster band patterns across samples. The pre-processing involves peak detection and batch effect corrections. In particular, we propose a local scoring algorithm for peak detection that is computationally efficient and performs well for minor peaks (Section 2.2). The detected peaks then enter the image alignment method that corrects batch effects in two steps: reference alignment and spatial dewarping. First, reference alignment calibrates multiple gels towards a common molecular standard. We perform piecewise linear stretching/compression by placing knots at the *marker* or *reference* bands present on all the images (Section 2.3.1). The reference-aligned gel images produce a set of peak locations that are then fitted by a novel hierarchical Bayesian model. The proposed model assumes that the smooth spatial gel deformations have deviated the observed peaks from their true landmarks. We use Markov chain Monte Carlo to estimate both the smooth warping functions and, for each detected peak, the posterior probabilities over a grid of landmarks where it is aligned. The Bayesian framework has the advantage of incorporating inherent uncertainty in assigning a peak to a molecular weight landmark.

The *aligned* high-frequency intensity data (Section 3.2) may be the input of many methods including hierarchical clustering, latent class models and factor analyses. In this paper, we focus on illustrating the value of alignment for the standard hierarchical clustering applied to data with known and unknown clusters (Section 4). At each iteration of the MCMC sampling, we obtain the multivariate binary signatures that represent autoantibody presence or absence over a grid of landmarks and align the gel images. Upon hierarchically clustering the aligned intensities at each iteration, we obtain a collection of dendrograms. In particular, we use the standard correlation-distance based agglomerative hierarchical clustering to create nested subgroups. For *N* samples, hierarchical clustering produces a dendrogram that represents a nested set of clusters. Depending on where the dendrogram is cut, between 1 and *N* clusters result. We demonstrate through real data that pre-processing more clearly separates the estimated clusters and improves the accuracy of cluster detection compared to naive analyses done without alignment.

The rest of the paper is organized as follows. Section 2 introduces the importance of pre-processing GEA data followed by algorithmic details for peak detection in Section 2.2 and batch effect correction in Section 2.3. In Section 3, we describe model posterior inference by MCMC and the statistical property of the shrinkage priors. We demonstrate how the proposed methods function through an application to signature estimation and subgroup identification of scleroderma patients in Section 4. The paper concludes with a discussion on model advantages and opportunities for extensions.

## 2. Data Pre-Processing

### 2.1 GEA Data and Pre-processing Overview

Gel electophoresis for autoantibodies (GEA) is designed to separate autoantibody mixtures according to molecular weight and to radioactively map them as bands along the gel. Figure 1(a) shows an example of a raw GEA image. We tested four sets of samples from scleroderma patients with a malignancy; of note, these sera were pre-selected as being negative for the three most commonly found scleroderma autoantibodies (anti-topoisomerase 1, anti-centromere and anti-RNA polymerase III antibodies, which in aggregate are found in about 60% of scleroderma patients).

**Figure 1.**
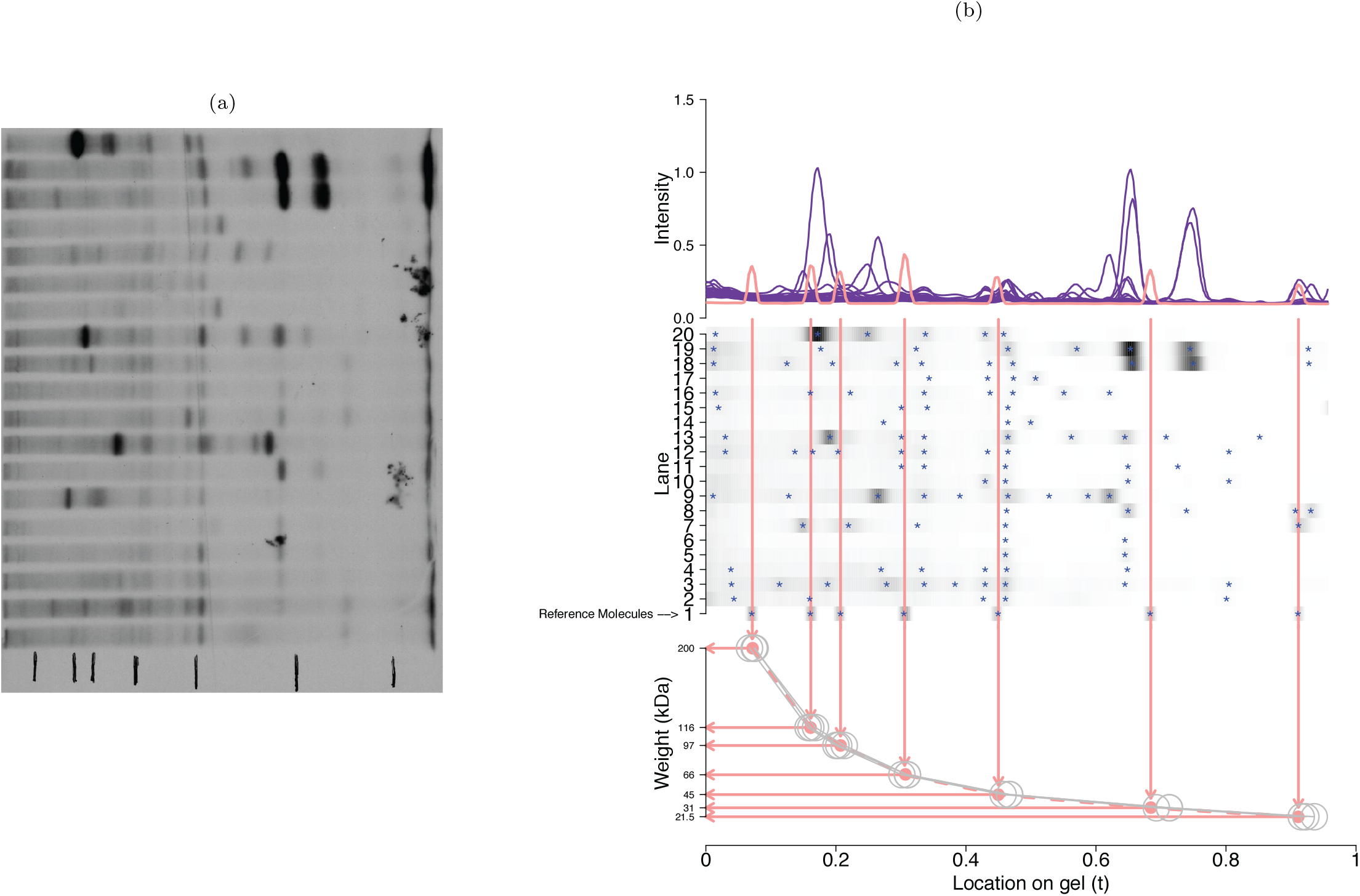
Gel electrophoresis autoradiography data for 20 samples on one gel. a) Raw GEA image. b) *Top*: Radioactive intensities for all the samples; *Middle*: Heatmap of the radioactive intensities for all the samples. The blue asterisks (*) denote the detected peaks. Seven vertical red lines indicate the locations of the seven reference molecules observed on lane 1. *Bottom*: Actual molecular weights (Y-axis) as read from the location along the gel (X-axis). Four location-to-weight curves are shown here, each corresponding to reference lane 1s in the four gels analyzed in Section 4.3 (the dashed red curve “- - -” is for the gel shown in the middle). Note the reference molecule misalignment shown by the scattered “O”.

Each sample set consisted of 19 patient sera plus one reference. In the middle panel of Figure 1(b), seven red vertical lines indicate the reference molecules of known weight (200, 116, 97, 66, 45, 31, 21.5) kDa. It also shows the band patterns that read out autoantibodies present in each of 19 patient samples (lanes 2-20). The top of Figure 1(b) shows the intensities from all the lanes; Seven clear spikes above the vertical lines again correspond to the reference molecules. The bottom of Figure 1(b) shows the piecewise linear interpolation of the location-to-weight function using the seven reference weights as knots. The weight of an arbitrary peak (“*”) can then be read from this interpolation. For example, the protein actin (about 42kDa) produced the peaks immediately to the right of the 45kDa reference (the fifth vertical line from the left). The misalignment of the actin peaks is caused by non-rigid image deformation (Section 2.3.2).

Identical reference molecules fail to align (empty circles, bottom panel of Figure 1(b)) across multiple gels because of variation in experimental conditions such as the strength of the electric field. We correct such misalignment by matching the reference peak locations across gels and then piecewise-linearly stretch or compress each gel using the reference peaks as knots. The technique is referred to as *piecewise linear dewarping* and was first used in human motion alignment anchored at body joints (e.g., Uchida and Sakoe, 2001).

The autoradiographic process is also vulnerable to smooth non-rigid gel deformation. This is most evident from the bands of actin, a ubiquitous protein of molecular weight 42 kDa, present in all lanes at around 0.43 (middle panel of Figure 1(b)). The bands form a smooth curve from the top to the bottom. The curvature represents the gel deformation since actin has identical weight and should appear at identical locations across the 19 lanes. Without correction, this deformation interferes with accurate sample comparisons even on the same gel. In Section 2.3.2, we propose a Bayesian hierarchical image dewarping model with shrinkage priors to correct the deformation and align the actin peaks.

To establish notations, let 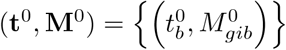 represent the standardized, high-frequency GEA data, for bin *b* = 1,*…, B* on lane *i* = 1,*…, N_g_* from gel *g* = 1,*…, G*. Appendix S1 describes the standardization of raw data. Here **t**^0^ is a equi-spaced grid over the unit interval [0, 1], where 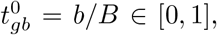 *b* = 1,*…, B*. 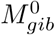 is the radioactive intensity scanned at 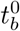 for lane *i* = 1,*…, N_g_*, gel *g* = 1,*…, G*. Let *N* = ∑*_g_ N_g_* be the total sample size.

For the rest of this section, we take the high-frequency data (**t**^0^, **M**^0^) and map it to multivariate binary data **Y** on a coarser common grid across gels. In Section 2.2, we propose a general method to transform an arbitrary high frequency, nearly continuous intensity data into raw peak locations. We first apply the peak detection algorithm to (**t**^0^, **M**^0^) and obtain the peak locations 𝒫^0^. In Section 2.3.1 we use the reference peaks, a subset in 𝒫^0^ from the first lane on each gel, to process (**t**^0^, **M**^0^) into reference-aligned data (**t**^R^, **M**^R^). In Section 2.3.2, we transform the peaks *P* detected from (**t**^R^, **M**^R^) to a joint posterior distribution of a *N* by *L* binary matrix **Y** that represents presence or absence of a peak over a grid of *L* landmarks for all the *N* samples (*L* = 100 in our application). In Section 3.2, we will process the reference-aligned high-frequency data (**t**^R^, **M**^R^) into (**t**, **M**) where the peaks appear at the landmarks indicated by the ones in **Y**.

### 2.2 Peak Detection

This section presents a general algorithm for detecting peaks from intensity data. We illustrate the algorithm by detecting peaks 𝒫^0^ from data (**t**^0^, **M**^0^). The peaks may appear with varying background intensities. Because the occurrence of a local maximum is thought to be more important than the background level in autoantibody signature estimation, we design the algorithm to be insensitive to the absolute intensity level.

We adopted the following peak detection algorithm:

i. *Local Difference Scoring*. For each bin *b* = 1,*…, B*, lane *i* = 1,*…, N_g_* of gel *g* = 1,*…, G*, calculate the local difference score by comparing the intensity at bin *b* to its left and right neighbors exactly *h* bins away and to the local minimum for locations in between (*t−h, t*+*h*) (truncated at 1 or *B* if *b* is near the endpoints). That is, we calculate

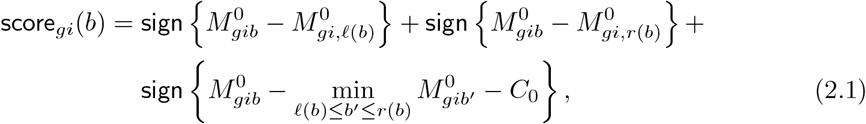

where sign(a) = 1, 0, *−*1 indicates positive, zero, or negative values; *l*(*b*) = max{*b − h*, 1} and *r*(*b*) = min{*b* + *h, B*} denote the left and right neighbors *h*(= 10) bins away, and *C*_0_ denotes the minimum peak elevation. The tunning parameter *h* controls the locality of the peaks and *C*_0_ controls the minimum peak magnitude.
ii. *Peak Calling*. We look for the bins among peak candidates defined by {*b |* score_*gi*_(*b*) = 3} that maximize their respective local intensities (see Appendix S2 for details and alternative peak calling methods). Let 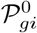 represent the collection of the peak locations for lane *i* and gel *g*.

#### Remark 1

The score defined in (2.1) depends only on the *sign*s of the differences in local intensities. They can be computed in parallel across all the samples. A two-dimensional analogue has been used in astrophysics to find low grey-scale intensity galaxies from telescope images (Xu *and others*, 2016).

### 2.3 Batch Effect Correction

#### 2.3.1 Reference Alignment via Piecewise Linear Dewarping

Molecules with identical weight do not appear exactly at the same location in each lane of a single gel due to gel deformation or across gels due to variations in experimental conditions. We first align the reference peaks 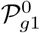, *g* = 1,*…, G* via piecewise linear dewarping to address the gel-to-gel variation (Uchida and Sakoe, 2001). In our application, we used seven reference molecules of known weight (200, 116, 97, 66, 45, 31, 21.5) kDa.

We first match the reference peaks 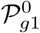 on a query gel *g* to the reference peaks 𝒫_*g*01_, on the template gel *g*_0_, and then use the matched reference peaks and the endpoints as knots to linearly stretch or compress the gels. Quadratic or higher-order dewarping is also possible, but we found linear dewarping performs sufficiently well for our data. Appendix S3 gives the details of the algorithm. We denote the high frequency, reference aligned data by 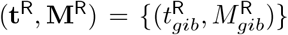. Applying the peak detection algorithm in Section 2.2 to this data, we collect all the detected peaks in 𝒫 = {𝒫*_g_, g* = 1,*…, G*} where 𝒫_*g*_ represents the peaks from gel *g*.

#### 2.3.2 Bayesian Image Dewarping to Correct Gel Deformation

Another source of error during autoradiographic visualization is the non-rigid, spatial gel deformation. The middle panel of Figure 1(b) shows one such example. It also reveals three analytical challenges to be addressed before obtaining meaningful results from an automatic disease subsetting algorithm. First, some proteins, e.g., actin, are detected on multiple gels and must be aligned. The blue asterisks that denote the detected peaks near 0.43, form a smooth but non-linear curve from the top to the bottom of the gel. Second, fewer bands appear on the right half of the image, because these smaller proteins tend to contain fewer methionine residues for radiolabeling. Higher estimation uncertainty of the dewarping function is therefore expected for the right half. Third, the observed locations of the peaks are likely random around their true locations as the result of the multiple sources of error.

To address these issues, we designed a hierarchical Bayesian dewarping algorithm for two-dimensional images. The algorithm simulates presence/absence data from the conditional distribution of protein occurrence over a grid of equispaced landmarks given the detected peaks 𝒫 from the prior preprocessing. The stochastic model is defined on a coarser grid of landmark proteins, ***ν*** = {0 = *ν*_0_ *< ν*_1_ *<* …*< ν_L_ < ν_L_*_+1_ = 1} where *ν_l_* = *l/*(*L* + 1), *l* = 0, 1,*…, L* + 1. In this paper, we will align peaks only to the internal knots {*ν_l_, l* ≠ 0, *L* + 1}; *ν*_0_ and *ν_L_*_+1_ will be used in the boundary constraint (2.5) to ensure endpoint alignment for all the sample lanes. We introduce a novel shrinkage prior to promote alignment of peaks to a common landmark. We also introduce shrinkage priors that regularize the overall smoothness of the spatial dewarping functions.

Let (*T_gij_, u_gij_*) denote the (location, lane number) for peak *j* = 1,*…, J_gi_* on lane *i* = 1,*…, N_g_*, gel *g* = 1,*…, G*. We fix *u_gij_* to take values in {1, 2,*…, N_g_* } and collect them in ***u*** = {*u_gij_* } where *u_gij_* = *u_gi_* if they belong to the same lane *i*. Let *P_g_* = ∑*_i_ J_gi_* denote the total number of peaks on gel *g* and *P* = ∑ *P_g_*. Let **T** = {***T**_g_*}, where ***T**_g_* = (…, *T_gi_*_1_, *T_gi_*_2_,*…, T_gi,Jgi_, …*)ʹ collects the peak locations for gel *g* = 1,*…, G*. Both ***u*** and **T** are *P*-dimensional column vectors. For computational stability, without changing notation, we standardize **T**, ***u*** and ***ν*** by substract their means and dividing by their standard deviations. We now use 𝒫 = {**T**, ***u***} to denote the data for the Bayesian dewarping model.

##### Model Likelihood

*Peak-to-landmark indicators* **Z**. Let *Z_gij_* take values in {1,*…, L*}. For example, *Z_gij_* = 3 indicates that the *j*-th peak in lane *i* on gel *g* is aligned to landmark 3. Let **Z** = {***Z**_g_, g* = 1,*…, G*} where ***Z**_g_* = {*Z_gij_, j* = 1,*…, J_gi_, i* = 1,*…, N_g_* }. Note that any **Z** can be converted to *N* multivariate binary observations **Y** = {(*Y_gil_, l* = 1,*…, L*)} for the presence or absence of a landmark, where *Y_gil_* = **1** {*l ∈* {*Z_gij_, j* = 1,*…, J_gi_*}}, referred to as *signature*.

*Gaussian mixture model for aligning observed peaks* **T**. We model **T** as observations from a Gaussian mixture model with *L* components, each representing one landmark. Given **Z** = {*Z_gij_* } and the spatial dewarping function 𝒮_*g*_ to be discussed later, we assume

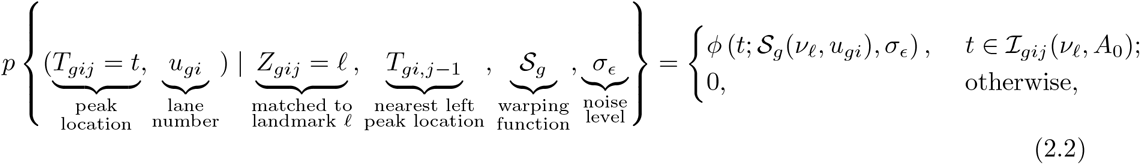

*l* = 1,*…, L*, for peak *j* = 1,*…, J_gi_*, lane *i* = 1,*…, N_g_*, gel *g* = 1,*…, G*, where *φ*(*·*; *a, b*) is the Gaussian density function with mean *a* and standard deviation *b*, and 𝒮*_g_* is an unknown smooth bivariate function that characterizes the deformation (*ν, u*) *↦* (𝒮_*g*_ (*ν, u*), *u*).

###### Remark 2

The peak location *T_gij_* is assumed to follow a Gaussian distribution with mean equal to *ν_l_* plus a horizontal displacement 𝒮_*g*_ (*ν_l_, u_gi_*) and noise level equal to *σ_∈_*. We assume *σ_∈_* is independent of landmark and lane. The density function (2.2) is positive only in the set *I_gij_* (*ν_l_, A*_0_) ≜ {*t*: *|t – ν_l_|< A*_0_ and *t > T_gi,j−_*_1_}. The first inequality prohibits *T_gij_* being matched to distant landmarks and limits the search space for *Z_gij_* in our algorithm; the second inequality places order constraints on the observed peak locations *T_gij_ > T_gi,j−_*_1_, *j* = 2,*…, J_gi_* − 1. We will restrict *Z_gij_ > Z_gi,j−_*_1_ to avoid reverse dewarping.

*Bivariate smooth warping functions 𝒮_g_*. For gel *g*, we model the warping function 𝒮_*g*_: ℝ^2^ → ℝ using the tensor product basis expansion

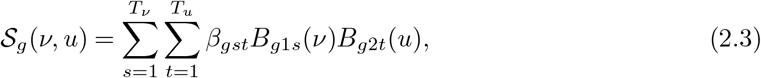

where *B_g_*_1*s*_(*·*) and *B_g_*_2*t*_(*·*) are the *s*-th and *t*-th cubic B-spline basis with intercept, and ***κ**_ν_* and ***κ**_u_* are the knots along the two coordinate directions, respectively (Friedman *and others*, 2001, Chapter 5) and *T_ν_* and *T_u_* are the total number of bases. In subsequent analyses, we choose ***κ**_ν_* with *T_ν_* − 4 internal knots at the *s/*(*T_ν_* − 3)-th quantile of {*T_gij_* }, *s* = 1,*…, T_ν_* − 4 and similarly for ***κ**_u_*. Let the two sets of B-spline basis functions along *ν*- and *u−* direction be ***B**_g_*_1_(*·*) = (*B_g_*_11_(*·*),…, *B_g_*_1*Tν*_ (*·*))ʹ and ***B**_g_*_2_(*·*) = (*B_g_*_21_(*·*),…, *B_g_*_2*Tu*_ (*·*))ʹ, respectively.

However, valid spatial gel deformations are limited to gel stretching, compression or shift along the *ν* direction. We thus constrain the shape of 𝒮*_g_, g* = 1,*…, G* by

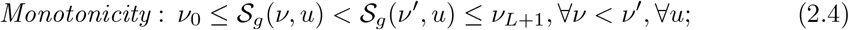

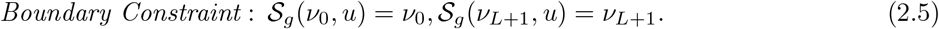

The first constraint prevents reverse gel dewarping and the second assumes no gel shifting. It can be relaxed to allow horizontal shifts by adding/substracting ∆ for both equalities. We implement both constraints by requiring the B-spline coefficients ***β**_g_* = {*β_gst_*} to satisfy: *ν*_0_ = *β_g_*_1_*_t_ < β_g_*_2*t*_ *<* …*< β_g,Tν −_*_1,*t*_ *< β_gTν t_* = *ν_L_*_+1_, ∀*t* = 1,*…, T_u_*. Although only sufficient for 𝒮_*g*_’s monotonicity and boundary constraints, the foregoing ***β**_g_* constraints allow flexible and realistic warpings. Figure 2 shows a member warping function that corrects for local “*L*”-, “*S*”- and “7”-shaped deformations.

**Figure 2.**
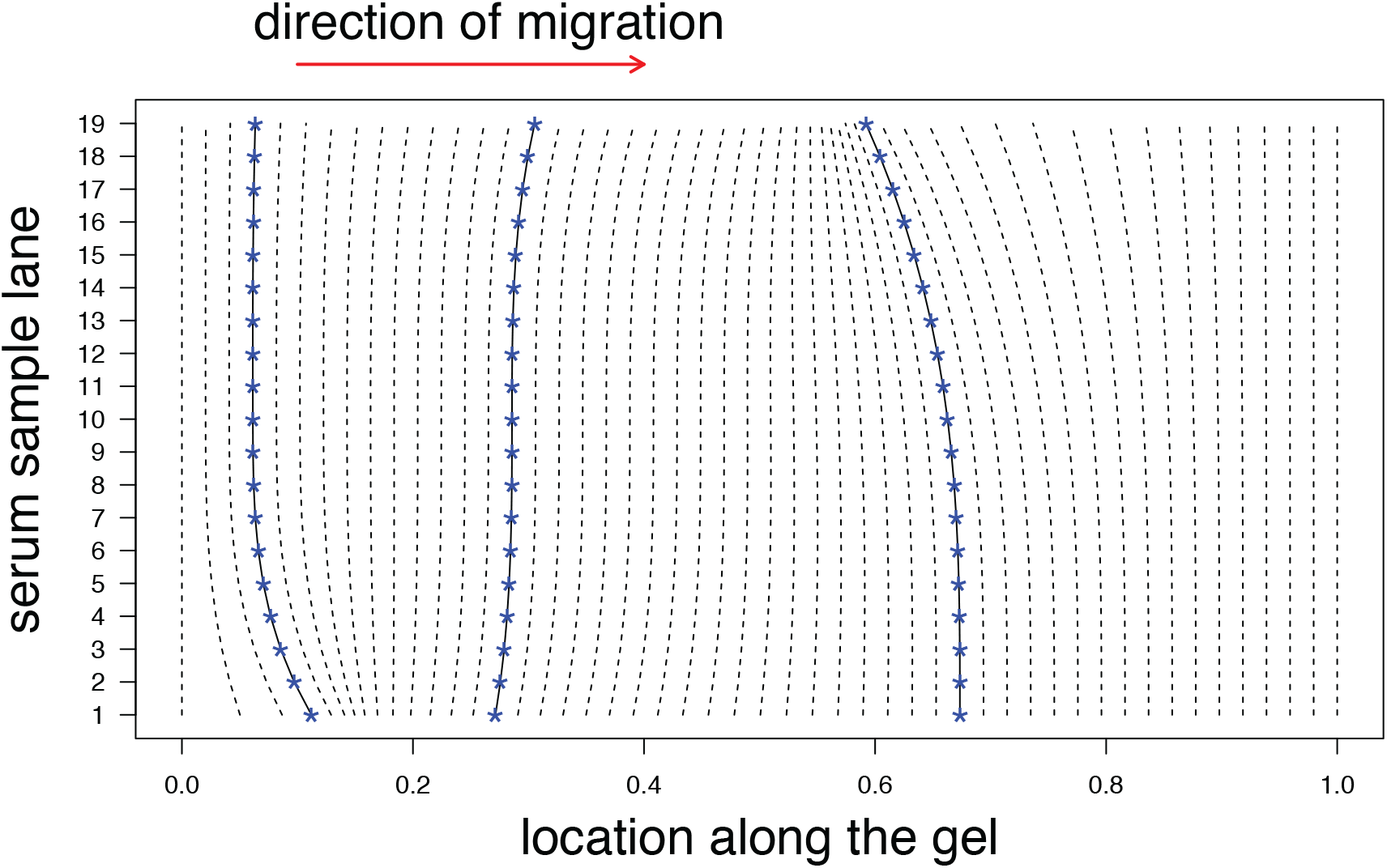
The posterior mean estimate of a gel warping function 𝒮 that corrects local stretching or compression. Highlighted are three vertical smooth curves, each of which aligns the peaks (blue asterisks “*∗*”) with identical molecular weights.

The likelihood function (2.2) models the misaligned data 𝒫 = {**T**, ***u***} in terms of the unknown spatial transformation 𝒮_*g*_ and the alignment **Z**. Multiple raw gel images are then aligned by the model estimates accompanied by model-based uncertainty quantification. Importantly, coherent image registrations must align the universal actin peaks and hence require the borrowing of information among multiple misaligned observations. We accomplish this by sharing a set of intensity parameters {*λ_l_*} among the gels.

##### Prior

*Prior for* **Z**. We describe a shrinkage prior for **Z** motivated by the need 1) to align the actin peaks (middle panel, Figure 1(b)), and 2) to share the information about the location of actin peaks across multiple gels.

We specify the prior distribution based on a discretized, non-homogeneous Poisson process with extreme intensities at a small number of landmarks. Let the total number of the observed peaks follow a Poisson distribution: 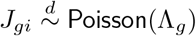, for sample *i* = 1,*…, N*, gel *g* = 1,*…, G*. Given *J_gi_*, let 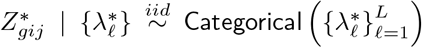 describe which landmarks are present in sample *i* of gel *g*. For sample lane *i*, we then define {*Z_gij_*} as the increasingly sorted 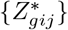. That is, we impose the order restriction *Z_gij_ ≤ Z_gijʹ_* whenever peak *jʹ* appears to the left of peak *jʹ* (*T_gij_ ≤ Tgij_ʹ_*). For hyperpriors, let 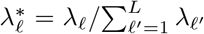 where 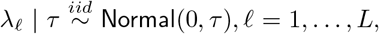, and the hyperparameter 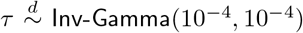. Integrating over *τ*, we obtain a marginal *t*-distribution for *λ_l_*.

###### Remark 3

It is easy to calculate the prior probability of landmark *l* present in a sample 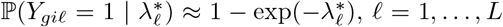, for large *L*. As shown by (A4) in Appendix, the ratio of the conditional posterior probabilities of assigning the peak *T_gij_* to landmark *l* versus *lʹ* is factorized into 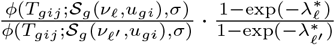. Suppose landmark *l* is associated with a higher intensity, i.e., 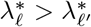, the second ratio favors landmark *l* given the likelihood ratio in the first term. Because the 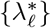 are independent of *g* and *i*, they globally modulate the probability of a landmark being present in all the gels. In our application, all the landmarks are *in a priori* assumed to be equally likely by specifying independent *t*-distributed priors for the *λ_l_*_s_. The *t*- distributions are heavy-tailed and can occasionally generate a large value of *λ_l_*_0_. Given *λ_l_*_0_, the posterior sampling algorithm will visit and then retain any configuration of **Z** that results in a large value of ∑*_g,i_ Y*_*gil*0_ if the configuration substantially increases the joint posterior.

***Prior for β_g_***. We incorporate the prior knowledge that large and abrupt image deformations are rare. We first specify priors for the horizontal basis coefficients *β_gst_*, *s* = 2,*…, T_ν_* − 1 at the *u*-direction basis *t* = 1. We use a first-order random walk prior (Lang and Brezger, 2004)

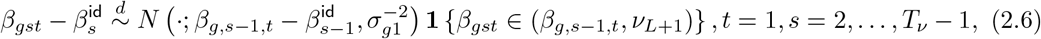

where 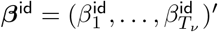 is the vector of coefficients to represent an identity function *I*: *ν ↦ ν* in terms of the bases 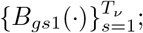 The truncation of *β_gst_* is needed for monotonicity (2.4). The hyperparameter 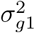 controls the similarity between 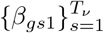 and ***β***^id^ and hence the similarity between 𝒮_*g*_ (*·, u*) and the identity function *I*; 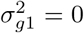 represents no warping. We refer to 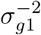 as the smoothing parameter along the *ν*-direction.

Next, for any *s* = 2,*…, T_ν_* − 1, we specify another random walk prior for the vertical basis coefficients

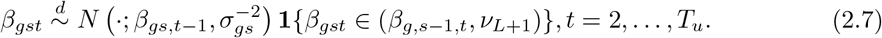

Similarly, the hyperparameter 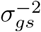 controls the smoothness of 𝒮_*g*_ along the vertical or *u*− direction; 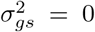 produces identical amounts of warping for all the lanes. Details about the hyperpriors for 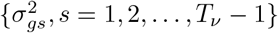 are provided in Appendix S4.

*Joint Distribution*. The joint distribution of all the unknowns is

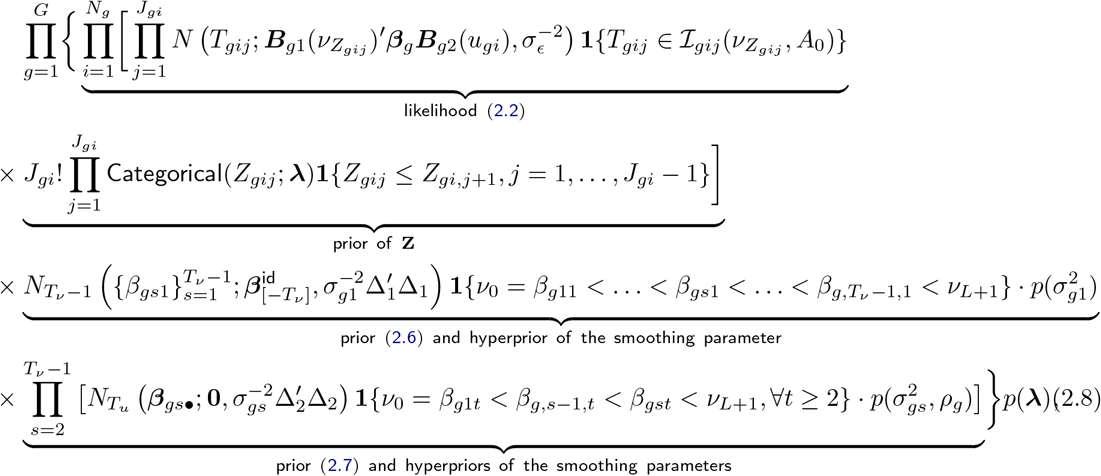

where *p*(***λ***), 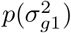 and 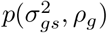 are the priors and hyperpriors and *N_d_*(*·*; ***µ***, **Λ**) denotes the *d*-dimensional multivariate normal density with mean ***µ*** and precision matrix **Λ** (can be degenerate). The matrix ∆_1_ maps a column vector to its first-order differences (used in 2.6): ∆_1*kk*_*1* = *δ*(*k* + 1, *k*ʹ) *− δ*(*k, k*ʹ), *k* = 1,*…, T_ν_* − 2, *k*ʹ = 1,*…, T_ν_* − 1, where *δ*(*a, b*) = 1 if *a* = *b* and equals 0 otherwise; Similarly we define ∆_2_ with *T_ν_* replaced by *T_u_* + 1.

## 3. Model Estimation and Implementation

### 3.1 Posterior Sampling

We use Markov chain Monte Carlo (MCMC) to simulate samples from the joint posterior distribution of all the unknowns (e.g., Gelfand and Smith, 1990) and then draw posterior inferences about chosen functionals of the model parameters. Of special interest are the gel warping functions {𝒮_*g*_ (*·, ·*; ***β***)} and the peak-to-landmark alignment **Z**. Appendix S5 describes the sampling algorithm and discusses conditions for statistical identifiability of the warping functions. All the model estimation and visualization are performed by the R package spotgear (https://github.com/zhenkewu/spotgear).

Turning to dewarping a new GEA image *g^∗^*, we perform reference alignment (Section 2.3.1) and then obtain the peaks *P_g_*∗ (Section 2.2). We approximate the joint posterior of (***β**_g_∗, **Z**_g_*∗) by

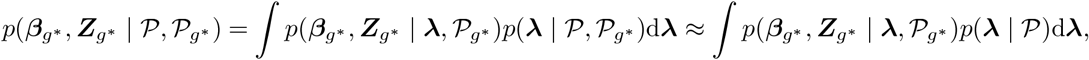

where the first term of the integrand is an one-sample conditional posterior and the second term is the posterior of ***λ*** given the old peaks 𝒫. Given 𝒫*_g_*∗, the first term can be derived from the joint distribution (2.8) with *G* = 1. The integral can then be approximated by *K^−^*^1^ ∑*_k_ p*(***β**_g_∗, **Z**_g_∗ |_**λ**_*(*k*), 𝒫*_g_*∗) where {***λ***^(*k*)^, *k* = 1,*…, K*} are the stored posterior samples.

### 3.2 Exact Peak Alignment for M

We now describe the high-frequency data (**t**, **M**) = (**t**, **M**(**Z**)) that have peaks *exactly* aligned according to **Z**. We simply perform piecewise linear dewarping for each sample lane so that the detected peak (*T_gij_, u_gi_*) is horizontally adjusted to match its landmark *ν_Zgij_*, *j* = 1,*…, J_gi_*. To do so, we apply the algorithm in Appendix S3 with {*ν*_0_, *T_gij_,…, T_giJgi_, ν_L_*_+1_} as a query and {*ν*_0_, *ν_Zgi_*_1_,*…, ν_ZgiJgi_, ν_L_*_+1_} as a template. In the following analyses, we will create aligned data using either 1) **Z** = **Z**^(*k*)^, *k* = 1,*…, K*, the stored MCMC samples when calculating the posterior distributions of the parameters that are functions of **Z**, or 2) **Z** = **Z**:= {*Ẑ_gij_* = arg max_*l*=1,…,*L*_ *p*(*Z_gij_* = *l | P*)}, the *maximum a posteriori* (MAP) alignment to obtain **M**(Ẑ).

#### Remark 4.

Because 𝒮_*g*_ is monotonic in *ν* given *u*, let 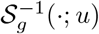 denote its inverse. One might be tempted to dewarp the images so that (*t, u*) is horizontally aligned to 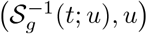. However, because a peak *T_gij_* varies around its mean *S_g_* (*ν_gij;ugi_*) unless 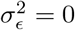, the inverse mapping cannot guarantee that 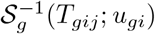 is equal to *ν_Zgij_*.

## 4. Applications to Scleroderma Patient Subsetting

Our methodology is motivated by the long-term clinical objective of finding an autoantibody signature that subsets autoimmune disease patients into groups with more homogeneous phenotypes and disease trajectories. The first step is to use the GEA data to cluster patients into subgroups with potential to have different outcomes. We used sera from well-characterized patients with scleroderma and an associated cancer identified through the IRB-approved Johns Hopkins Scleroderma Center database (Shah *and others*, 2017). To test our algorithms, we first analyze two GEA replicates each of 20 samples. Compared to the results of hierarchical clustering without pre-processing, we show our pre-processing method creates more clearly separated clusters. We also show our pre-processing improves the accuracy of cluster detection evaluated against the true matching. As a second test, we apply the pre-processing method to GEA measurements on 76 patients with unknown clustering. We observe that the use of the pre-proecessing algorithm identifies clusters that are clearly separated and scientifically meaningful.

### 4.1 Outline of Analyses

For subsequent analyses, this section describes the steps of pre-processing, clustering and three metrics that evaluate the obtained clusters.

#### Pre-processing

We apply the peak detection algorithm in Section 2.2 followed by batch effect corrections as described in Section 2.3. We exclude the reference lane on each gel when performing the two dimensional Bayesian dewarping. We used *T_ν_* = 10 and *T_u_* = 6 cubic B-spline basis functions in the horizontal and vertical directions, respectively. The dewarping functions are estimated by {𝒮̂_*g*_ = 𝒮_*g*_ (*·, ·*; β̂_*g*_)} where β̂_*g*_ is the posterior mean estimated by the empirical average of the MCMC samples. We also obtain the MAP estimate **Ẑ** = {Ẑ_*gij*_ }. The choice of the number of bases is crucial for the estimation of the warping functions (e.g., Lang and Brezger, 2004). For example, larger values of (*T_ν_, T_u_*) define a richer class of functions that can accommodate abrupt local image deformations. Visual inspection of the alignment of the actin peaks makes clear that more parsimonious models are preferred. Further improvements in knot selection is possible using knots on a nonequidistant grid so that more knots are placed where the spatial image warping is severe and the peaks are dense.

#### Clusterings

Given the peak-to-landmark alignment **Z**, we follow Section 3.2 to obtain peak-aligned images **M** = **M**(**Z**) and then obtain clustering solutions. For example, let **M** = **M**(Ẑ) where Ẑ is the MAP alignment. For each pair of sample *i* and *iʹ*, we calculate the pairwise distances *d*(*i, iʹ*) = 1 − cor(**M***_gi·_*, **M**_*giʹ*_) where **M**_*gi·*_ = (*M_gi_*_1_,*…, M_giB_*) *ʹ* and cor(*·, ·*) is the Pearson’s correlation coefficient. Denote the *N* by *N* matrix of pairwise distance s by D̂ = {*d*(*i, iʹ*)}. We use D̂ in the standard agglomerative hierarchical clustering with complete linkage to produce a dendrogram *Τ̂* = *Τ* (D̂). By varying the level of cutting the dendrogram *Τ̂*, we obtain a nested set of clusterings 𝒞̂(*n*), *n* = 2,*…, N*. We similarly denote the dendrogram produced without pre-processing by *Τ̂* ^0^ = *Τ* (*D*^0^) where *D*^0^ is the correlation-based distance matrix computed from **M**^0^. We denote the nested clusters by 𝒞̂^0^(*n*), *n* = 2,*…, N*. We will evaluate the obtained clusters by three criteria below.

#### Adjusted Rand Index

We assess the agreement between two clusterings of the identical set of observations using the adjusted Rand index (aRI; Hubert and Arabie (1985)). aRI is defined by

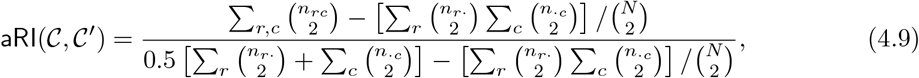

where *n_rc_* represents the number of observations placed in the *r*th cluster of the first partition *𝒞* and in the *c*th cluster of the second partition *𝒞*ʹ, 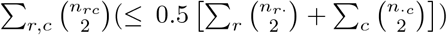 is the number of observation pairs placed in the same cluster in both partitions and 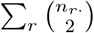 and 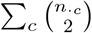 calculates the number of pairs placed in the same cluster for the first and the same cluster for second partition, respectively. aRI is bounded between −1 and 1 and corrects for chance agreement. It equals one for identical clusterings and is on average zero for two random partitions; larger values indicate better agreements between the two clustering methods.

#### Average silhouette

We also evaluate the strength of each clustering method using the *average silhouette* (Rousseeuw, 1987). For observation *i*, its silhouette *s*(*i*) for a partition *𝒞* compares the within- to the between-cluster average distances: *s*(*i*) = [*b*(*i*) *− a*(*i*)]*/*max{*a*(*i*), *b*(*i*)} where *a*(*i*) is the average distance of *i* to all other observations within the same cluster and *b*(*i*) = *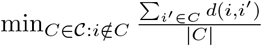* is the minimum average distance between *i* and a cluster not containing *i*. *s*(*i*) lies in [−1, 1] where a large value indicates observation *i* is in a tight and isolated cluster. A larger *average* silhouette *s*¯(*𝒞*) = *N^−^*^1^ ∑*_i_ s*(*i*) indicates more clearly separated and tighter clustering 𝒞.

#### Confidence levels of clusters

In addition to the alignment uncertainty addressed by the posterior distribution [**Z** *|* 𝒫], another source of uncertainty is the clustering of the aligned high-frequency intensity data (**t**, **M**(**Z**)) given **Z**. In this paper we chose not to specify the full probability distribution for the continuous intensities (**t**, **M**(**Z**)). Following Shimodaira *and others* (2004) and Efron *and others* (1996), we use bootstrap resampling to assess the confidence in the estimated dendrogram *Τ̂* (setting **Z** = **Ẑ**, the *MAP*). The bootstrap method perturbs the data by randomly sampling the columns of (**t**, **M**(**Ẑ**)) with replacement and assesses the confidence levels for the presence of each subtree in *Τ̂*. We calculate the frequency with which a subtree appears in an estimated dendrogram across all the bootstrap iterations where a large value (e.g., *>* 0.95) indicates strong evidence. We similarly bootstrap (**t**^0^, **M**^0^) to assess the confidence in the dendrogram *Τ̂*^0^ estimated without alignment.

### 4.2 Replication Experiments

Each of 20 biological samples were tested with two different lengths of exposure to autoradiographic devices: long (two-week) versus short (one-week) exposure. We ran 40 lanes on two gels that form 20 replicate pairs. Each gel image has 20 sample lanes: 19 serum sample lanes plus one reference lane comprised of molecules with known weights. The posterior dewarping results are shown in Appendix Figure S2.

We assess the agreement between the estimated clustering 𝒞̂^(*k*)^(*n*) and the true replication-based clusters 𝒞^∗^ by aRI(𝒞̂^(*k*)^(*n*), 𝒞^∗^), for the number of clusters *n* = 2,*…*, 20 and the stored MCMC iteration *k* = 1,*…, K*. At iteration *k*, 𝒞̂^(*k*)^(*n*) is the clustering solution obtained by cutting the dendrogram that hierarchically clusters the peak-matched data **M**(**Z**^(*k*)^) where **Z**^(*k*)^ is drawn from the posterior [**Z** *|* 𝒫].

The pre-processing enhances the hierarchical clustering to produce clusters closer to the true replicate pairs. In Figure 3, the posterior mean of the adjusted Rand indices based on *K* = 5, 000 saved MCMC samples (solid line, *K^−^*^1^ ∑_*k*_ aRI(𝒞̂^(*k*)^(*n*), 𝒞^∗^)) are uniformly higher than the adjusted Rand indices based on data without pre-processing (dashed line, aRI(𝒞̂0(*n*), 𝒞^∗^)). In the bottom panel, for every *n*, the posterior distribution for the difference between the two aRIs excludes zero increases with the numbers of clusters. In addition, the pairwise distances in D̂ (obtained from **M**(**Z**̂)) decreased by between 6.2 and 66.4% (mean 26.9%) relative to those in *D*^0^. These decreases in the distances result in a dendrogram *Τ̂* that puts 13 replicate pairs at the terminal leaves as compared to 8 in *Τ̂*^0^.

**Figure 3.**
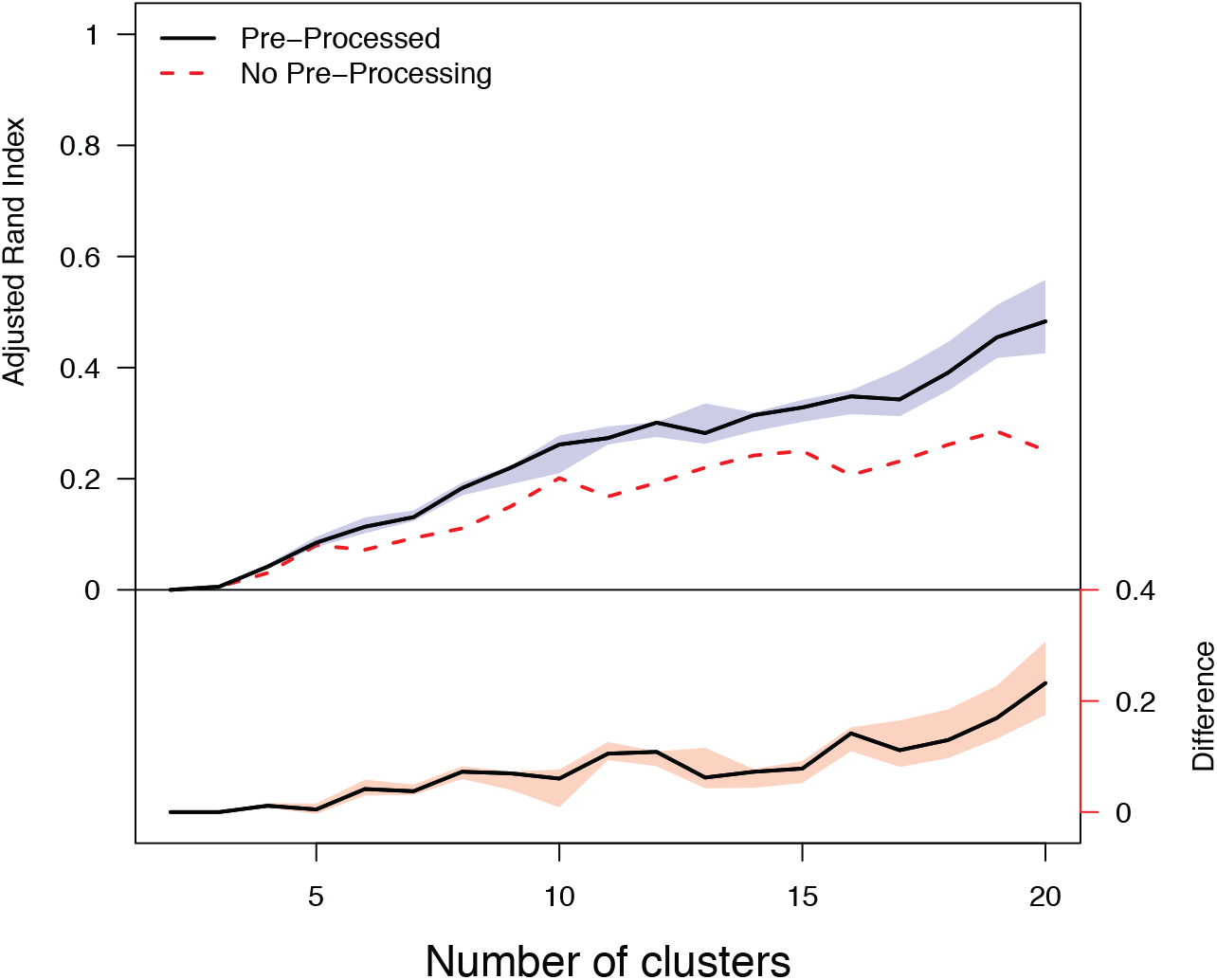
Comparison between the adjusted Rand Indices obtained with and without pre-processing. *Top*: The solid line and the blue shaded area represent the posterior mean aRIs and the pointwise 95% credible intervals, respectively. The dashed line is based on **M**^0^ without pre-processing. *Bottom*: The solid line represents the difference between the aRIs obtained with and without pre-processing: *K^−^*^1^ ∑_*k*_ aRI(𝒞̂^(*k*)^(*n*), 𝒞^∗^) aRI(𝒞̂^0^(*n*), 𝒞^∗^), *n* = 2,*…*, 20; the shared area shows the pointwise 95% credible intervals.

We also observed uniformly increased confidence levels of the presence of true replicate pairs upon pre-processing. Appendix Figure S3 examines the confidence levels associated with each sub-tree with (hierarchically clustering the MAP-aligned data (**t**, **M**(**Ẑ**)) and without pre-processing. For example, for pair 18, the estimated confidence level increases from 0.71 to 1 after pre-processing; The confidence levels for detecting the pairs 2, 8 and 11 see similar increases from 0.67, 0.79, 0.66 to 0.97, 0.86, 1, respectively. The increase in confidence levels is partly explained by the tighter clusters obtained after data pre-processing: the average silhouette computed from the MAP clustering, *s*¯ (𝒞̂(*n*), increased 14.2 − 117.6% (0.03 − 0.18 in magnitude) for *n* = 2,*…*, 20 clusters.

### 4.3 Scleroderma GEA Data without Replicates

We ran 4 GEA gels, each with 19 patient sera and one reference lane. The sera are from scleroderma patients with cancer who are all negative for common autoantibodies to RNA polymerase III, topoisomerase I and centromere proteins. We had no other prior knowledge about known or novel autoantibodies at the time the study was conducted. The sera were loaded in random order on each gel; the reference sample comprised of known molecules was always in the first lane. In the following, we describe the estimated dewarping, alignment and the resulting clusters.

#### Dewarping

We pre-process the four gel sets by estimating the dewarping functions {𝒮*_g_, g* = 1, 2, 3, 4} and the peak-to-landmark alignment **Z**. We first removed a few spots on the right of the gels caused by localized gel contamination and assumed absence of peaks at these spots. The posterior dewarping results are shown in Figure 4. Each detected peak {*Τ_gij_* } (blue dot) is connected to its matched MAP landmark Ẑ_*gij*_ (red triangle). The vertical bundle of black curves, one per landmark, visualizes the global shape of the estimated warping functions 𝒮̂_*g*_. Along each estimated vertical curve, the locations J(𝒮̂_*g*_ (*ν_R_, u*), *u, ∀u* represent identical molecular weights.

**Figure 4.**
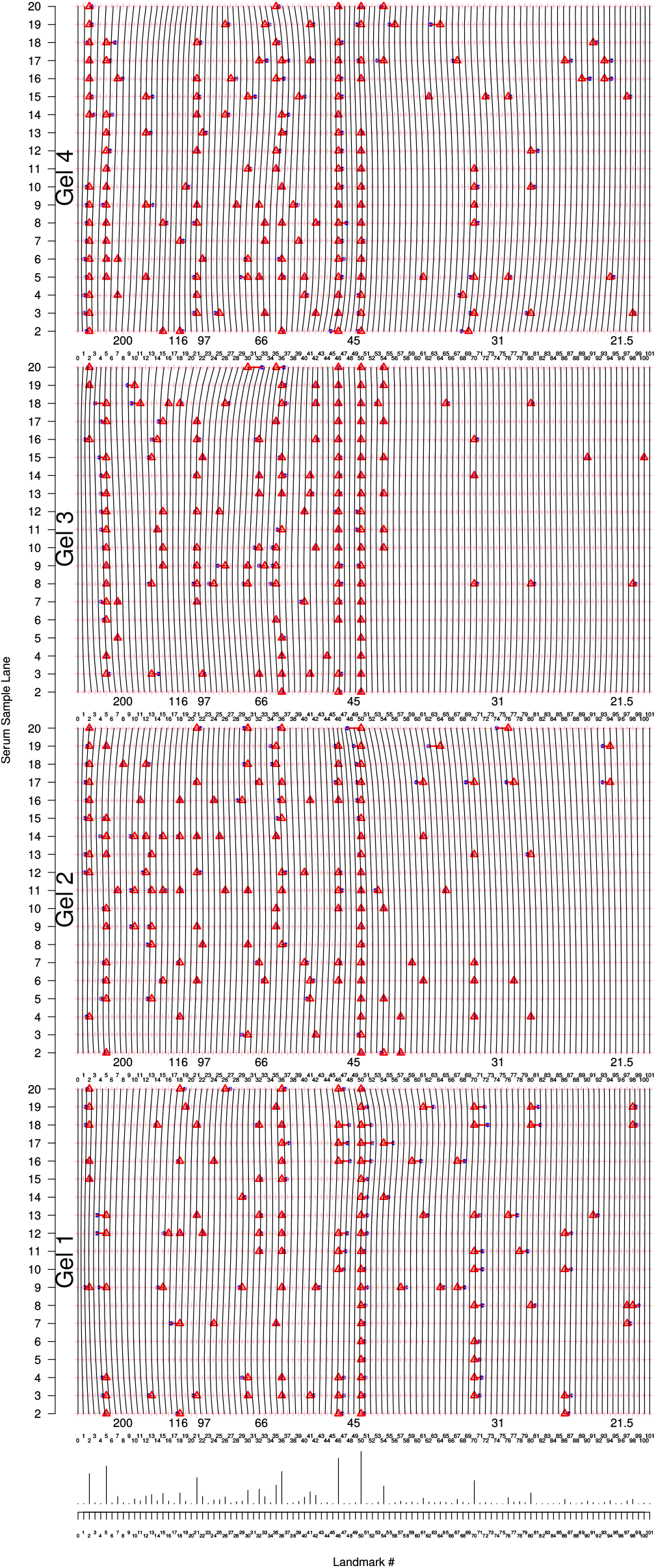
Bayesian spatial dewarping results for the experiment without replicates. *Top:* For each gel set, 19 serum lanes over a grid of *L* = 100 interior landmarks (reference lanes excluded). Each detected peak *T_gij_* (solid blue dots “*·*”) is connected to its *maximum a posteriori* landmark Ẑ_*gij*_ (red triangle “∆”). The image deformations are shown by the bundle of black vertical curves {*u* ↦ 𝒮_*g*_ (*ν_l_, u*): *u* = 2,*…*, 20, *l* = 1,…, *L, g* = 1, 2, 3, 4, each of which connects the estimated locations of identical molecular weight. *Bottom:* Marginal posterior probability of each landmark protein present in a sample.

#### Alignment to landmarks

The marginal posterior probabilites of each landmark in a sample are shown at the bottom of Figure 4. For example, the posterior probability is 0.59 for landmark 50 (about 43.4 kDa, actin): the MAP estimate Ẑ shows that 73 out of 76 lanes. The marginal posterior probability is expected to further increase when more samples containing actin are analyzed via hierarchical Bayesian dewarping. Landmark 46 (about 46.6 kDa) is another autoantibody hotspot where 54 out of 76 lanes have matched peaks. On the other hand, only 18 and 1 out of 76 are matched to Landmarks 36 (about 59.8 kDa) and 89 (about 23.4 kDa), respectively. Their marginal posterior probabilities are hence low at 0.21 and 0.01.

An animation of the continuous dewarping process is available at https://github.com/zhenkewu/spotgear. It matches the detected peaks *Τ_gij_* to their MAP landmarks *Z*̂_*gij*_ and morphs the posterior mean dewarping 𝒮̂_*g*_ into the constant function *I*: (*ν, u*) *↦* (*ν, u*). Also shown is the pre-processed high-frequency data (**t**, **M**(Ẑ)) with exactly matched peaks as described in Section 3.2.

#### Clusters

Our pre-processing method removed global warping phenomena and revealed a few strong clusters. The clusters with 0.95 confidence levels or higher are shown in red boxes in Figure 5 for the analyses done with pre-processing (top) and without pre-processing (bottom). A comparison of the two clustering solutions favors the pre-processing approach. For example, within the dendrogram at the top, the first cluster from the right (number 44) consists of seven sample lanes ((Set, Lane): (1,19), (4,3), (1,18), (3,8), (4,10), (2,4), (2,13)) that are enriched at roughly 32.7 and 27.9 kDa. This group is split into two clusters (numbers 47 and 14) for the analyses done without pre-processing. In a second example, the cluster 46 at the bottom and cluster 40 at the top are comprised of identical samples (enriched at about 103.4 kDa). We observe the confidence level increases from 0.97 to 1 after pre-processing. Pre-processing again produced more clearly separated clusters and eliminated many large clusters that are otherwise formed at the bottom of Figure 5; We observed 8.8 − 39.5% increases in the average silhouette based on the MAP alignment.

**Figure 5.**
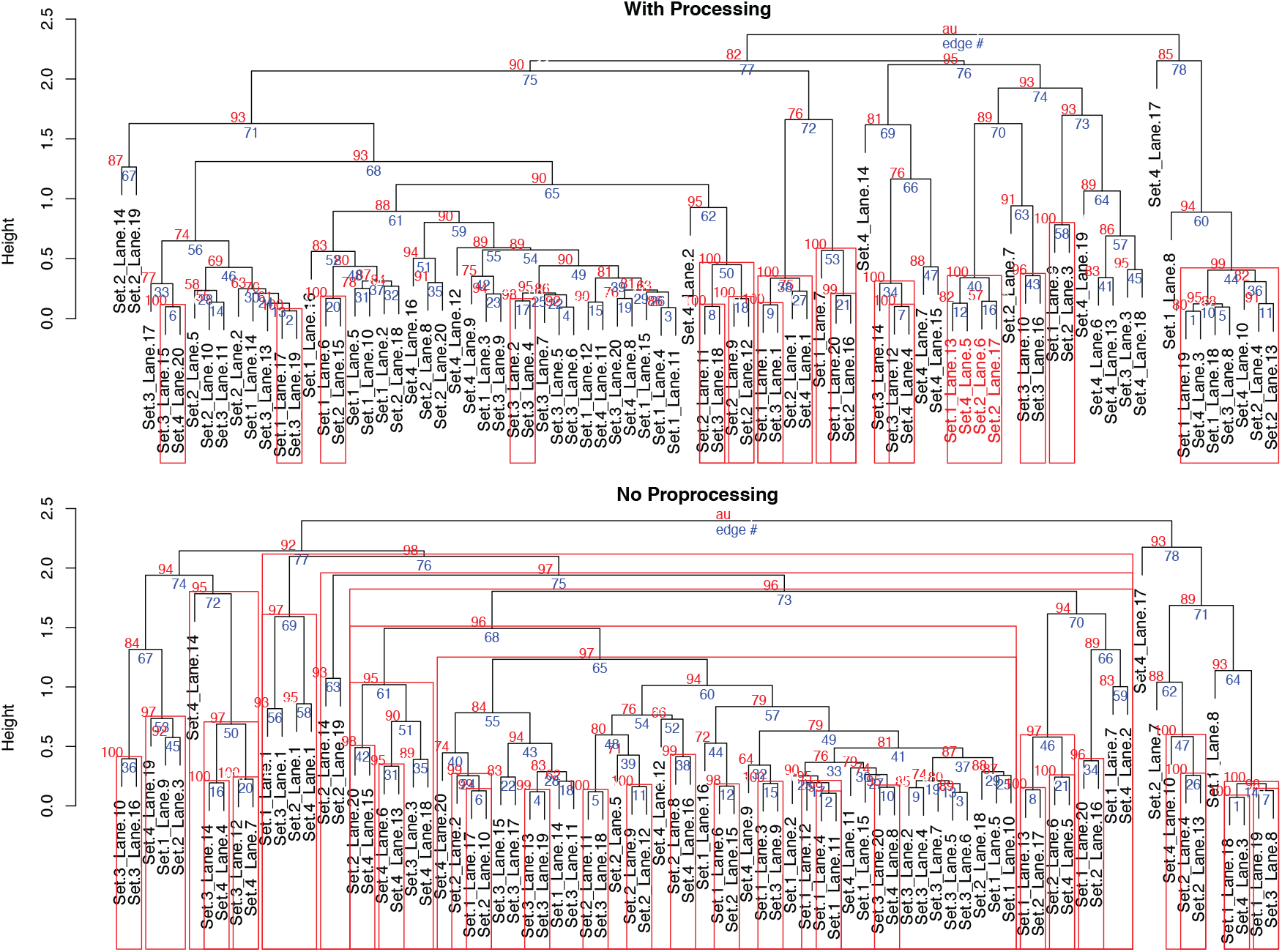
Estimated dendrograms with (top) and without (bottom) pre-processing for the second data set. The red boxes show the subtree appearing in *>* 95% of bootstrapped dendrograms with the actual estimated frequencies shown in red on top of the subtrees.

## Discussion

In this article, we have developed a novel statistical approach to pre-processing and analyzing two-dimensional image data obtained from gel electrophoresis autoradiography (GEA). Our objective is to eliminate artifactual data patterns that can confound our ability to use standard clustering algorithms such as hierarchical clustering to detect subsets of autoimmune disease patients. The hierarchical Bayesian image dewarping model provides a natural framework for assessing uncertainty in the estimated alignment and warping functions and allows us to make inferences about many functions of parameters.

In Section 4, we analyzed two sets of data from scleroderma patients. For the data with replication, we showed that the adjusted Rand indices increased if we perform pre-processing prior to standard hierarchical clustering. Based on the MAP alignment, the average silhouette that measures the strength of clustering increased by 14 to 118%. The pre-processing also increased the confidence levels for detecting true replicates.

For the data without replicates, we showed that our pre-processing method successfully aligned the actin peaks. It also increased the confidence levels for the clusters that appeared in both clusterings (one with pre-processing and the other without pre-processing).

We conclude that there is added benefits of applying the pre-processing procedure prior to estimating disease subsets. We expect marginal though worthwhile gains to be achievable by using more carefully designed and tested tuning parameter selection procedure for local scoring (Section 2.2).

In the analysis of data with out replicates (Section 4.3), we grouped the samples by creating a *single* dendrogram given a fixed **Z** = Ẑ, i.e., the MAP alignment. Uncertainty exists in both the alignment and the dendrogram obtained by hierarchical clustering. We have addressed the former by the posterior distribution [**Z** *|* 𝒫] and the latter by bootstraping. Future work is needed to assume a likelihood involving the unknown dendrogram structure to obtain and represent its posterior uncertainty (e.g., Chakerian and Holmes, 2012).

Two extensions based on prior biological knowledge are the current subject of further research. First, in our hierarchical Bayesian dewarping model, we assumed that the intensity parameters 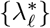 are shared among the samples. However, the prevalence of autoantibodies may differ by subpopulation. For example, cancer versus non-cancer patients may have distinct distributions for the abundance of certain autoantibodies. We can either add another hierarchy on top of 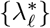 or develop regression models for 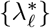 to incorporate disease phenotype information and covariates such as age and gender.

Second, proteins in the cells tend to work in complexes, so multiple autoantibodies are likely to be produced against a particular protein complex. This mechanism can be represented by a binary matrix **E**_*C×L*_ where the *c*-th row (*E_c_*_1_,*…, E_cL_*) is a multivariate binary vector with 1 for presence of landmark *f* in complex *c* and 0 otherwise. The complexes are then assembled via ***η**_N ×L_* = **AE** to produce the actual presence or absence of the landmarks for every patient, where **A** is a *N × C* binary matrix where each row represents the presence or absence of the *C* complexes. Prior biological knowledge can be readily implemented via constraints on **A** or **E**. For example, *A_i_*_1_ = 1 for all the samples acknowledges the universal presence of autoantibodies produced against complex 1, e.g., actin and likely others. **A** and **E** can be inferred from the alignment indicators **Y** and continuous intensities. We may use regularization or shrinkage priors in a Bayesian framework to encourage a few maximally different complexes (e.g., Broderick *and others*, 2013). One practical advantage of the Bayesian factorization approach lies in its convenient accommodation of repeated GEA on the same unknown sample by placing equality constraints on the rows of **A**. Finally, our latent variable formulation ***η*** = **AE** makes it easy to incorporate multiple sources of patient lab and phenotype data that inform ***η***, facilitate subgroup definition by **A** and perform individual predictions via the posterior predictive distributions of ***η*** (e.g., Coley *and others*, 2016; Wu *and others*, 2016, 2017).

## Supplementary Materials

Supplementary Material is available at the website of this paper.

## Acknowledgments

We thank the Editors, Associate Editor and two referees for their constructive suggestions that improved the presentation of our method and results. Research reported in this work was partially funded through a Patient-Centered Outcomes Research Institute (PCORI) Award (ME-1408-20318), NIH grant K23 AR061439 (to AS) and generous grants from the Jerome L. Greene Foundation and the Donald B. and Dorothy L. Stabler Foundation. The Johns Hopkins Rheumatic Disease Research Core Center, where the sera were processed and banked, and the antibody assays were performed, is supported by NIH grant P30 AR-070254.

## References

Broderick, Tamara, Jordan, Michael I. and Pitman, Jim. (2013, 08). Cluster and feature modeling from combinatorial stochastic processes. Statist. Sci. 28(3), 289–312.

Chakerian, John and Holmes, Susan. (2012). Computational tools for evaluating phylogenetic and hierarchical clustering trees. Journal of Computational and Graphical Statistics 21(3), 581–599.

Coley, Rebecca Yates, Fisher, Aaron J., Mamawala, Mufaddal, Carter, Herbert Ballentine, Pienta, Kenneth J. and Zeger, Scott L. (2016). A bayesian hierarchical model for prediction of latent health states from multiple data sources with application to active surveillance of prostate cancer. Biometrics, In Press.

Efron, Bradley, Halloran, Elizabeth and Holmes, Susan. (1996). Bootstrap confidence levels for phylogenetic trees. Proceedings of the National Academy of Sciences 93(23), 13429–13429.

Friedman, Jerome, Hastie, Trevor and Tibshirani, Robert. (2001). The elements of statistical learning, Volume 1. Springer Series in Statistics Springer, Berlin.

Gelfand, Alan E and Smith, Adrian FM. (1990). Sampling-based approaches to calculating marginal densities. Journal of the American statistical association 85(410), 398–409.

Hubert, Lawrence and Arabie, Phipps. (1985). Comparing partitions. Journal of classification 2(1), 193–218.

Lang, Stefan and Brezger, Andreas. (2004). Bayesian p-splines. Journal of computational and graphical statistics 13(1), 183–212.

Rosen, Antony and Casciola-Rosen, Livia. (2016). Autoantigens as partners in initiation and propagation of autoimmune rheumatic diseases. Annual review of immunology 34, 395–420.

Rousseeuw, Peter J. (1987). Silhouettes: a graphical aid to the interpretation and validation of cluster analysis. Journal of computational and applied mathematics 20, 53–65.

Shah, Ami A., Xu, George, Rosen, Antony, Hummers, Laura K., Wigley, Fredrick M., Elledge, Stephen J. and Casciola-Rosen, Livia. (2017). Brief report: Anticrnpc-3 antibodies as a marker of cancer-associated scleroderma. Arthritis & Rheumatology 69(6), 1306–1312.

Shimodaira, Hidetoshi *and others*. (2004). Approximately unbiased tests of regions using multistep-multiscale bootstrap resampling. The Annals of Statistics 32(6), 2616–2641.

Uchida, Seiichi and Sakoe, Hiroaki. (2001). Piecewise linear two-dimensional warping. Systems and Computers in Japan 32(12), 1–9.

Wu, Zhenke, Deloria-Knoll, Maria, Hammitt, Laura L and Zeger, Scott L. (2016). Partially latent class models for case–control studies of childhood pneumonia aetiology. Journal of the Royal Statistical Society: Series C (Applied Statistics) 65(1), 97–114.

Wu, Zhenke, Deloria-Knoll, Maria and Zeger, Scott L. (2017). Nested partially latent class models for dependent binary data; estimating disease etiology. Biostatistics 18(2), 200.

Xu, Bingxiao, Postman, Marc, Meneghetti, Massimo, Seitz, Stella, Zitrin, Adi, Merten, Julian, Maoz, Dani, Frye, Brenda, Umetsu, Keiichi, Zheng, Wei *and others*. (2016). The detection and statistics of giant arcs behind clash clusters. The Astrophysical Journal 817(2), 85.

